## Supplementary figures for "TIPE drives a cancer stem-like phenotype by promoting glycolysis via PKM2/HIF-1α axis in melanoma"

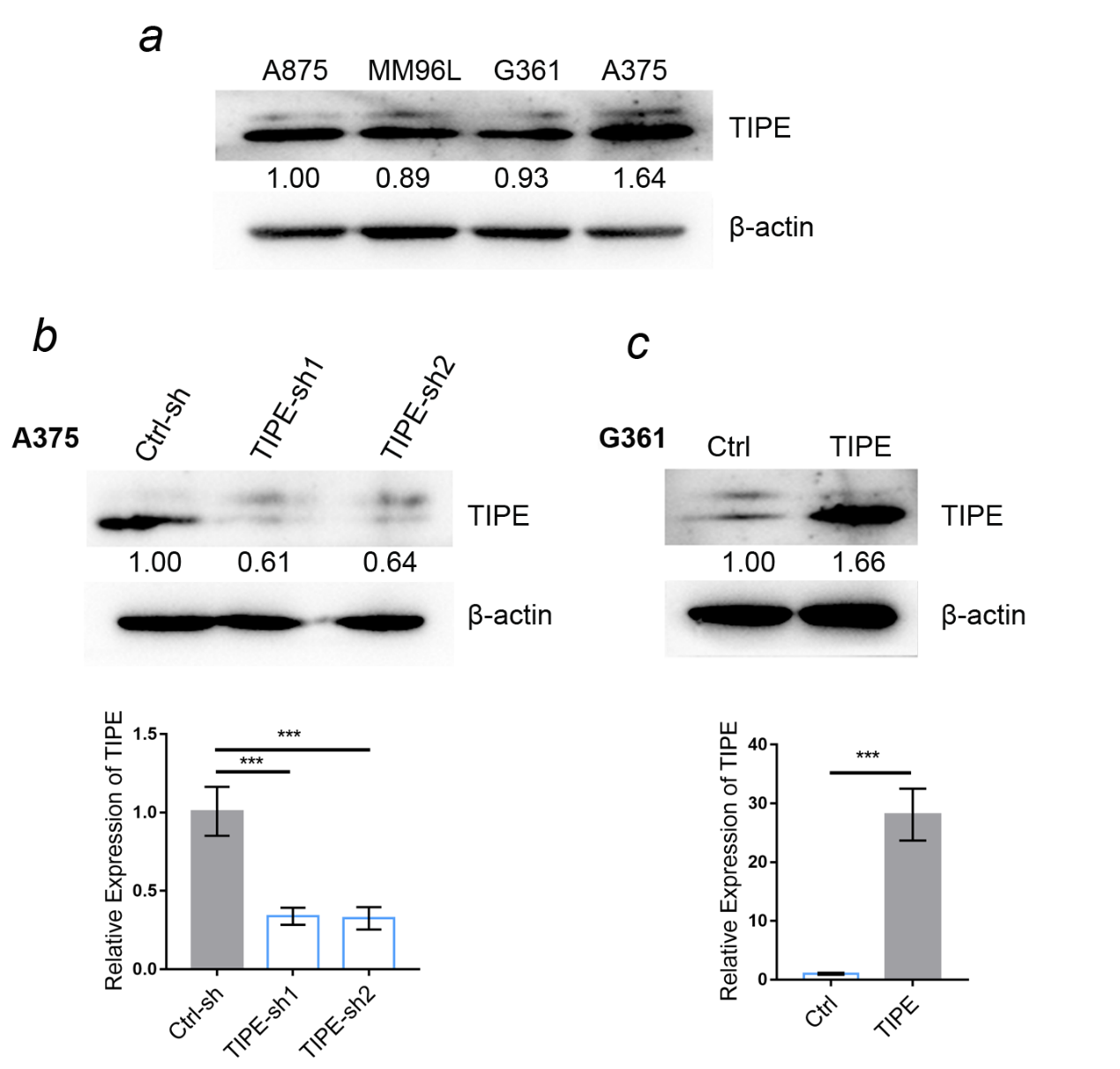

**Figure S1: *a*.** Western blot validation of TIPE in melanoma cell lines. ***b*.** Western blot and *q*PCR analysis of TIPE expression after TIPE interference in A375 cells. ***c*.** Western blot and *q*PCR analysis of TIPE expression after overexpression of TIPE in G361 cells.

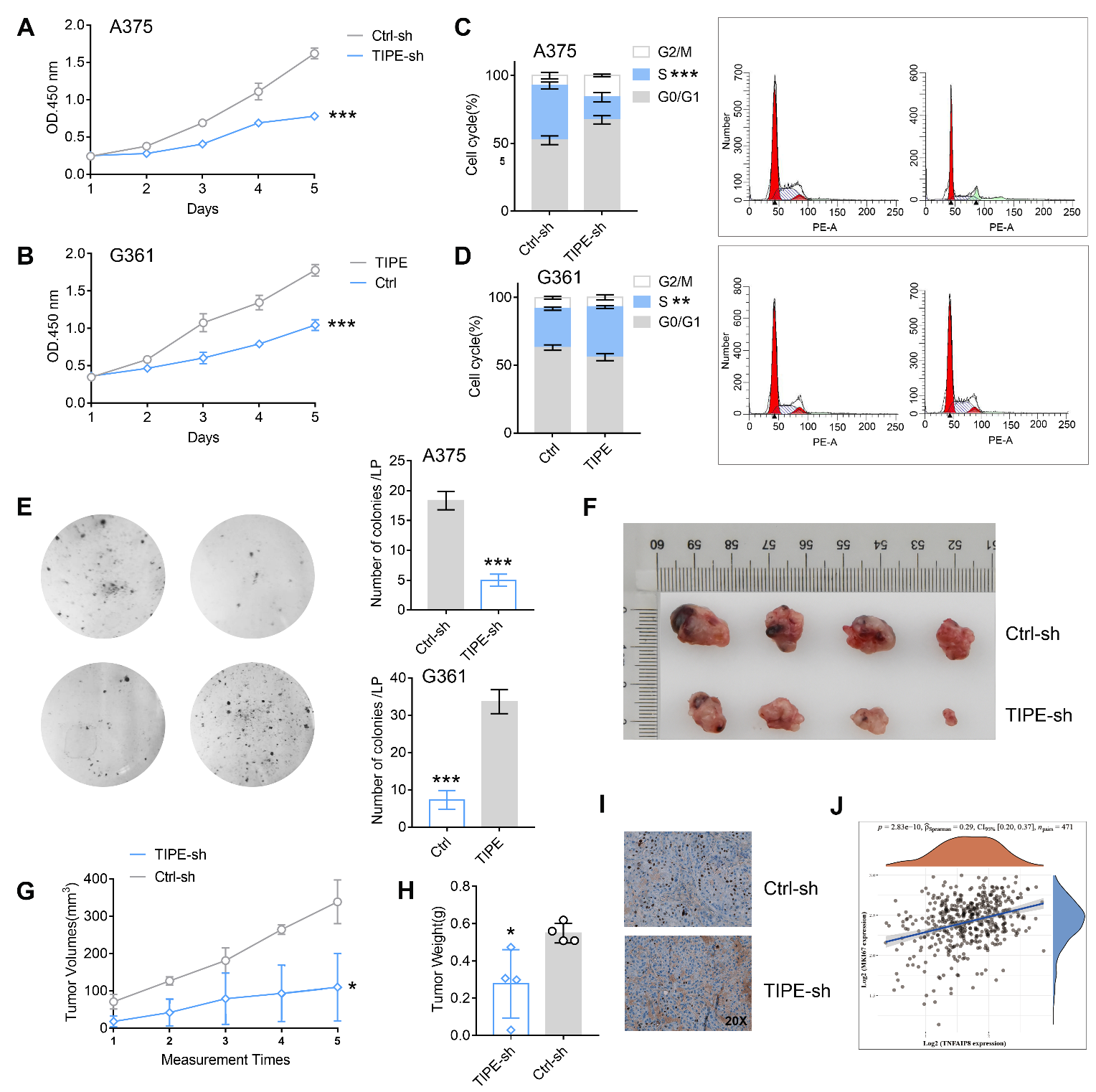

**Figure S2.** TIPE promotes melanoma cell proliferation in vitro and in vivo. ***a***. CCK8 assay showed that interfering TIPE (TIPE-sh) decreased the cells proliferation in A375 cells compared to control (Ctrl-sh). ***b***. Overexpression of TIPE(TIPE) elevated G361 cell proliferation compared to control (Ctrl). ***c, d***. FACS analysis of the effects of TIPE on cell cycle. ***e***. Effects of TIPE on colony formation in A375 and G361 cells. ***f-h***. Effect of TIPE on tumor formation in a nude mouse xenograft model. Nude mice were subcutaneously injection of cells containing Ctrl-sh and TIPE-sh, respectively. The tumor volume was measured every 7 days for 5 times until sacrifice. Representative images of tumors from the TIPE-sh and control groups (Ctrl-sh), n=4 for each group. ***i***. Interfering TIPE decreases the expression of Ki67 in nude mice, measured by immunohistochemistry. ***j***. TCGA dataset shows a positive relation between the mRNA expression of TIPE and Ki67 in melanoma patients. *P<0.05; **P<0.01; ***P<0.001.

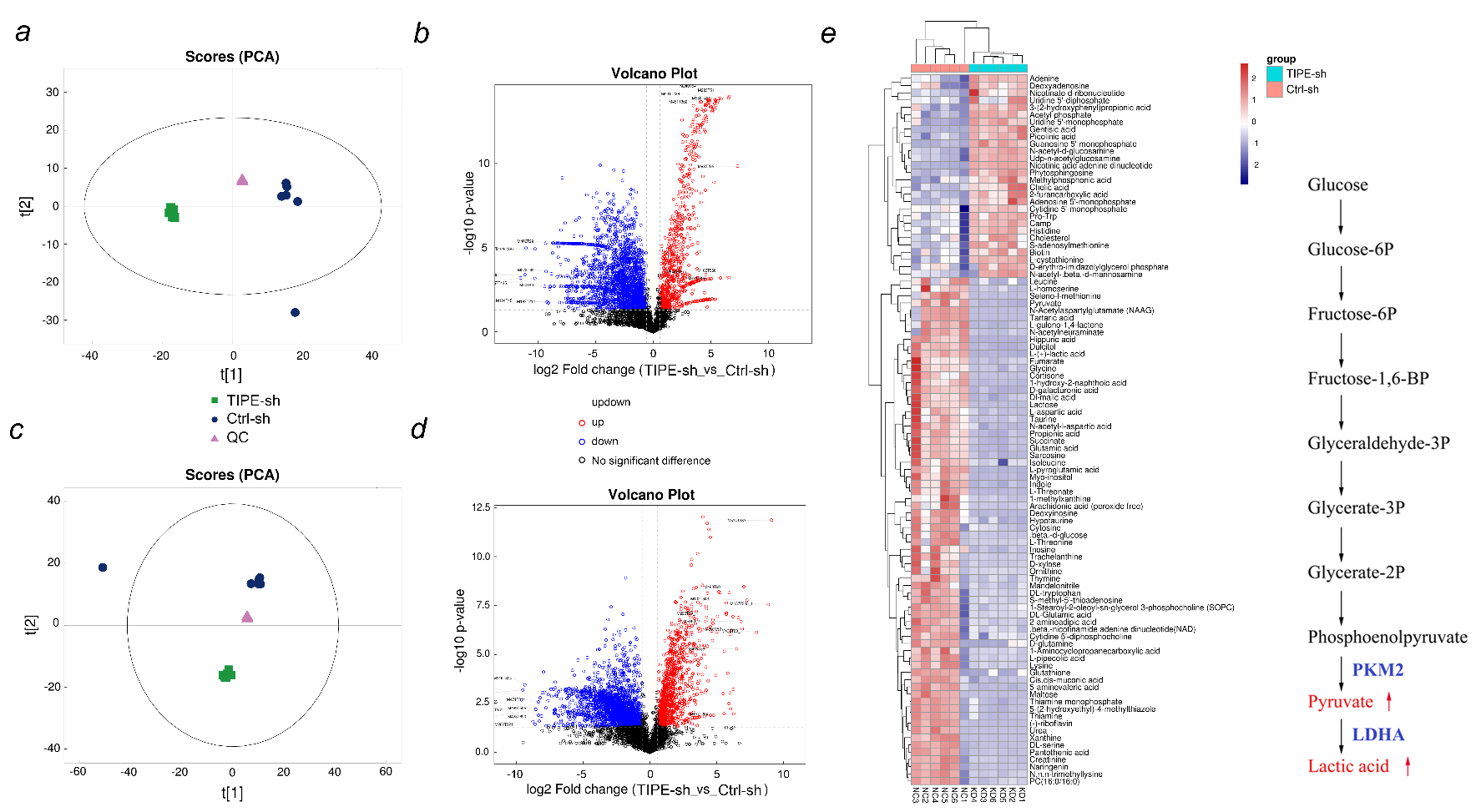

**Figure S3:** Principal component analysis (PCA) and volcano plots of the samples from TIPE interference group vs. control in the negative mode (***a, b***) or positive mode (***c, d***) by using untargeted metabolomics. ***e*.** Heatmap indicated that the glycolysis pathway including pyruvate and lactic acid is decreased after TIPE interference.

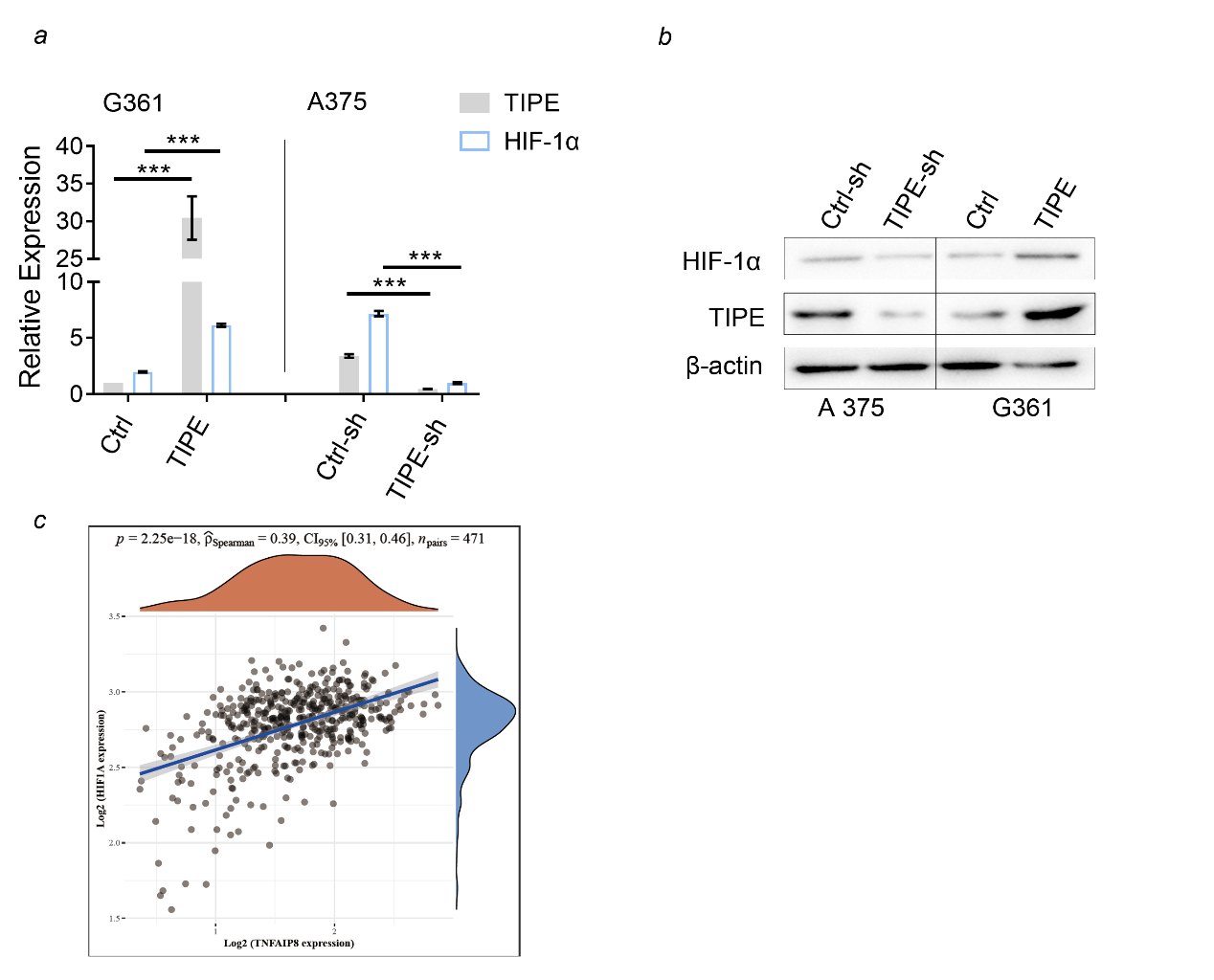

**Figure S4: *a*.** TIPE upregulated HIF-1α mRNA expression using a *q*PCR assay. ***b*.** Western blot analysis showed that TIPE increased HIF-1α expression. ***c*.** TCGA dataset indicated that TIPE has a positive correlation with HIF-1α.

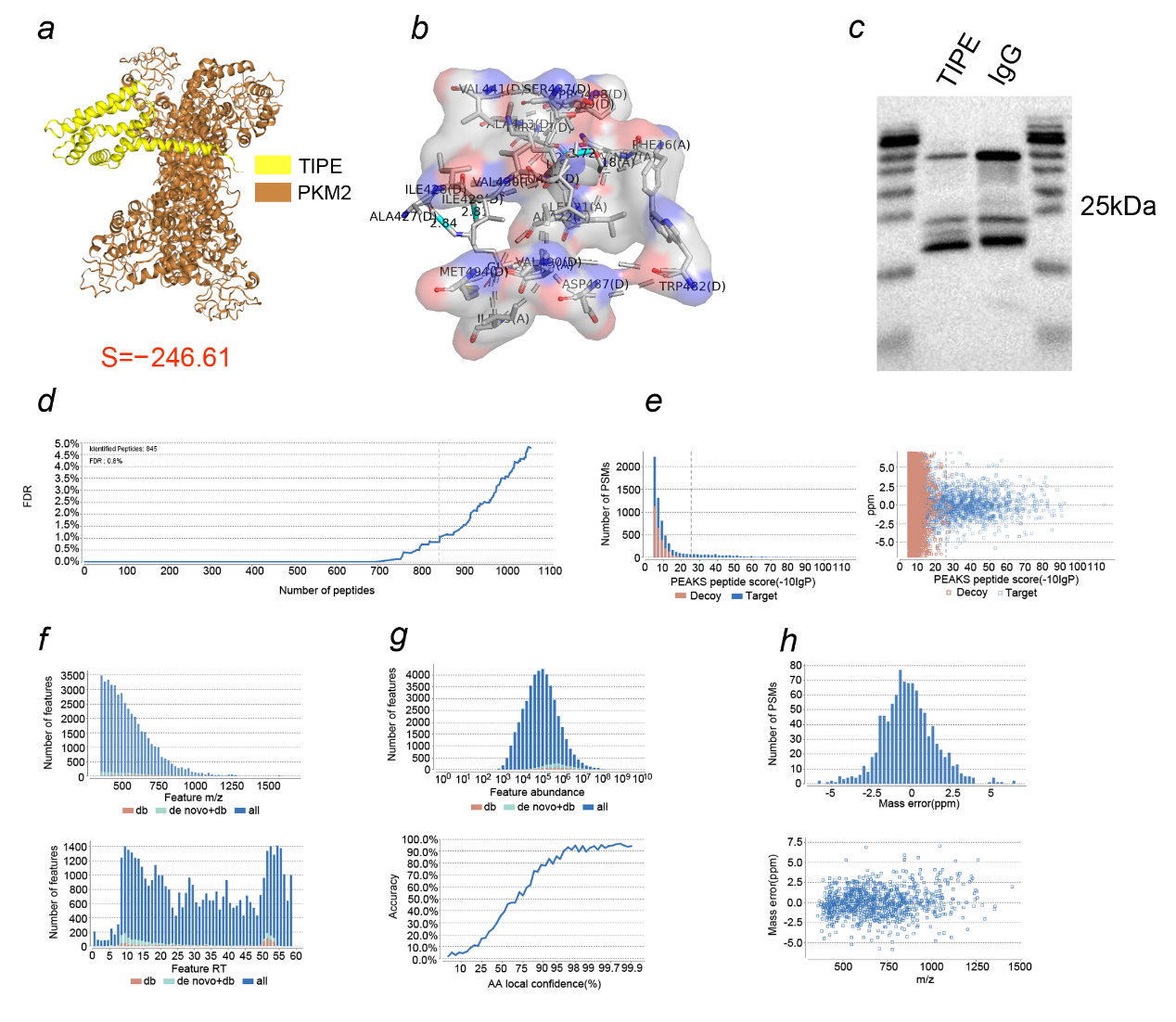

**Figure S5: *a, b*.** Molecular docking with a superposition of the three structures showed binding domain of TIPE and PKM2 by using PyMol 2.2.0(S=-246.61). ***c*.** Western blot analysis by using the TIPE antibody to bait its potential partner prior to the “IN-GEL DIGESTION” step. Before the “nano-HPLC-MS/MS ANALYSIS” step, the quality control including false discovery rate (***d***), score distribution (***e***), peptides feature detection (***f***),	verifiable de novo sequences (***g***) and quality accuracy control (***h***) were performed.

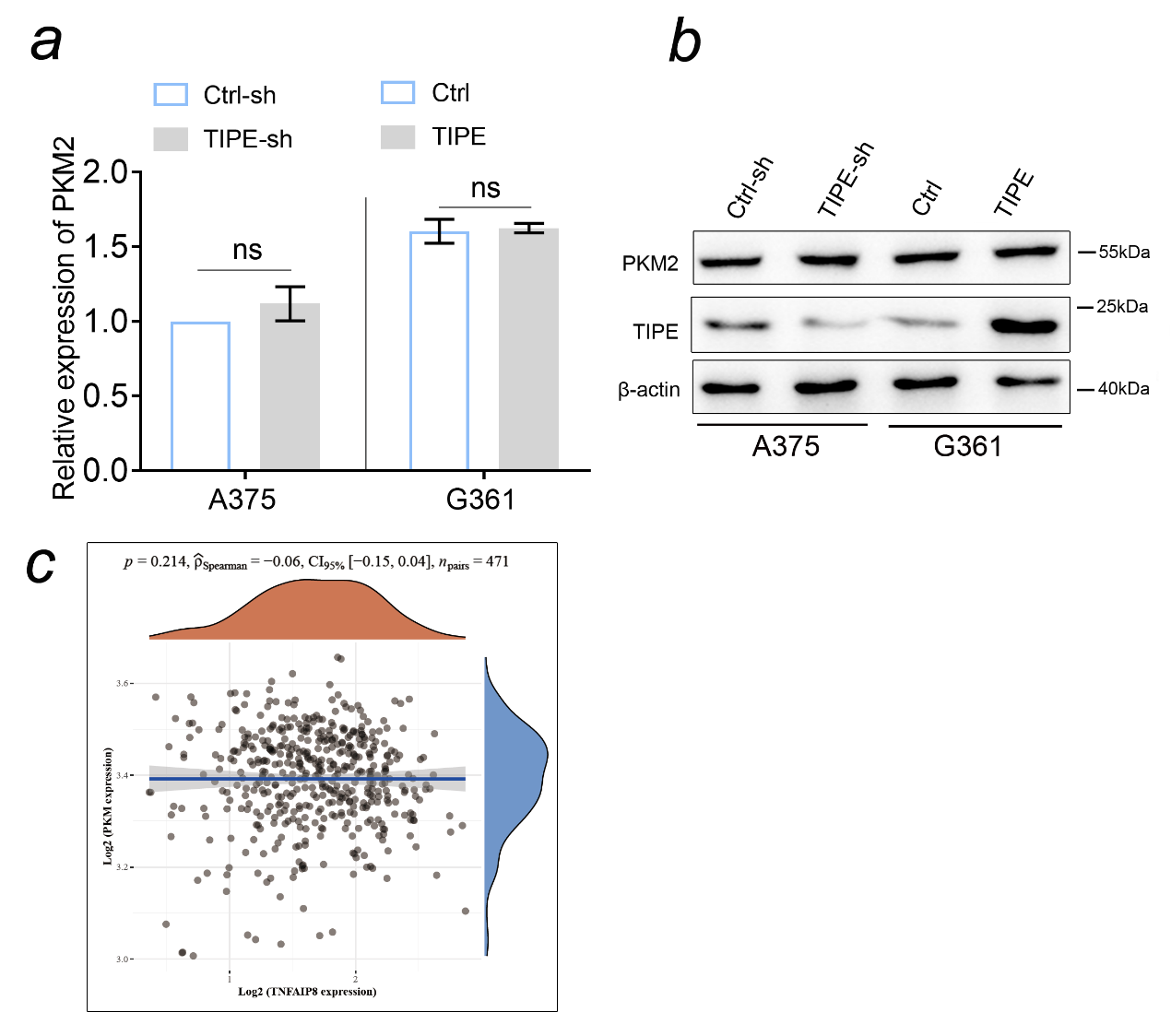

**Figure S6: *a,b*.** TIPE did not impact the expression levels of PKM2. ***c*.** TCGA dataset showed that there was no significant correlation between TIPE and PKM2.

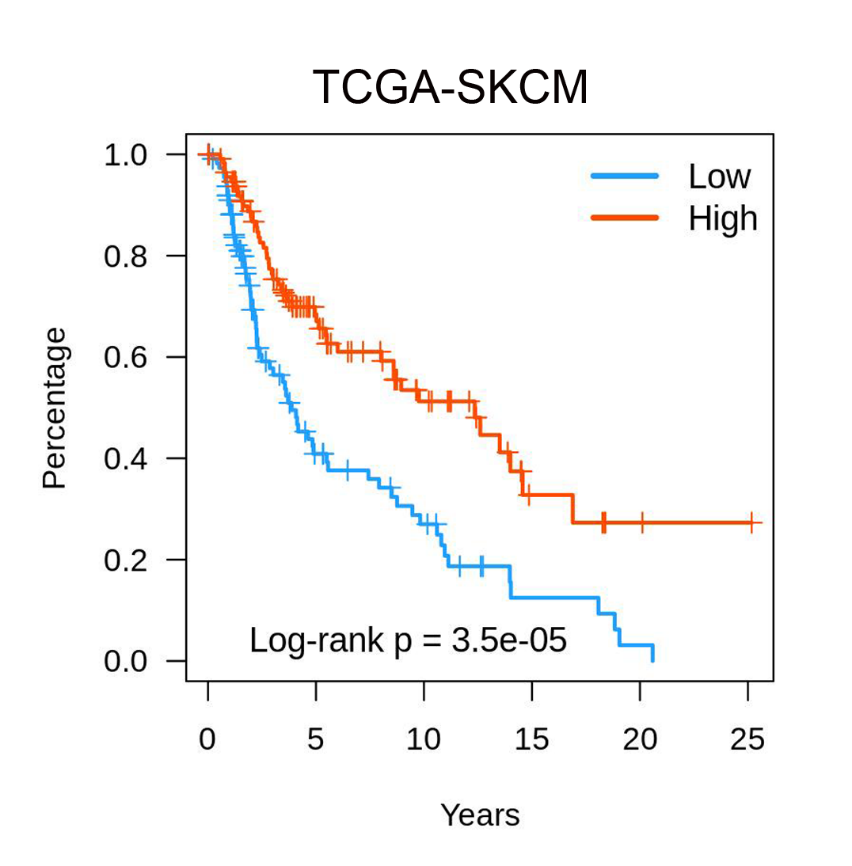

**Figure S7:** TCGA dataset showed a paradoxical observation that elevated TIPE expression is associated with a favorable prognosis in melanoma patients.
