## Supplementary tables for "TIPE drives a cancer stem-like phenotype by promoting glycolysis via PKM2/HIF-1α axis in melanoma"

**Table S1.** **The top candidate interacting proteins of TIPE identified by mass spectrometry**

| **Accession** | **Protein name** | **Gene name** | **Coverage(%)** | **#Peptides** |
| --- | --- | --- | --- | --- |
| P35527 | Keratin type I cytoskeletal 9 | KRT9 | 54 | 29 |
| P60709 | Actin cytoplasmic 1 | ACTB | 45 | 12 |
| P02751 | Fibronectin | FN1 | 11 | 22 |
| P68363 | Tubulin alpha-1B chain | TUBA1B | 37 | 10 |
| Q9BQE3 | Tubulin alpha-1C chain | TUBA1C | 33 | 9 |
| P13647 | Keratin type II cytoskeletal 5 | KRT5 | 34 | 24 |
| P35579 | Myosin-9 | MYH9 | 6 | 6 |
| P02533 | Keratin type I cytoskeletal 14 | KRT14 | 36 | 15 |
| P21333 | Filamin-A | FLNA | 5 | 6 |
| O60832 | H/ACA ribonucleoprotein complex subunit DKC1 | DKC1 | 17 | 11 |
| P63267 | Actin gamma-enteric smooth muscle | ACTG2 | 16 | 5 |
| P68032 | Actin alpha cardiac muscle 1 | ACTC1 | 16 | 5 |
| P68133 | Actin alpha skeletal muscle | ACTA1 | 16 | 5 |
| P62736 | Actin aortic smooth muscle | ACTA2 | 16 | 5 |
| Q14498 | RNA-binding protein 39 | RBM39 | 17 | 7 |
| P11387 | DNA topoisomerase 1 | TOP1 | 12 | 8 |
| P09211 | Glutathione S-transferase P | GSTP1 | 29 | 4 |
| P05787 | Keratin type II cytoskeletal 8 | KRT8 | 23 | 14 |
| P08779 | Keratin type I cytoskeletal 16 | KRT16 | 19 | 10 |
| **P14618** | **Pyruvate kinase PKM** | **PKM** | **19** | **5** |
| Q7Z794 | Keratin type II cytoskeletal 1b | KRT77 | 10 | 9 |
| P13646 | Keratin type I cytoskeletal 13 | KRT13 | 10 | 6 |

**Table S2. Genes up-regulated in response to low oxygen levels (hypoxia) in melanoma**

**Hypoxia score was calculated by single sample GSEA (ssGSEA)**

*ACKR3*

*ADM*

*ADORA2B*

*AK4*

*AKAP12*

*ALDOA*

*ALDOB*

*ALDOC*

*AMPD3*

*ANGPTL4*

*ANKZF1*

*ANXA2*

*ATF3*

*ATP7A*

*B3GALT6*

*B4GALNT2*

*BCAN*

*BCL2*

*BGN*

*BHLHE40*

*BNIP3L*

*BRS3*

*BTG1*

*CA12*

*CASP6*

*CAV1*

*CAVIN1*

*CAVIN3*

*CCN1*

*CCN2*

*CCN5*

*CCNG2*

*CDKN1A*

*CDKN1B*

*CDKN1C*

*CHST2*

*CHST3*

*CITED2*

*COL5A1*

*CP*

*CSRP2*

*CXCR4*

*DCN*

*DDIT3*

*DDIT4*

*DPYSL4*

*DTNA*

*DUSP1*

*EDN2*

*EFNA1*

*EFNA3*

*EGFR*

*ENO1*

*ENO2*

*ENO3*

*ERO1A*

*ERRFI1*

*ETS1*

*EXT1*

*F3*

*FAM162A*

*FBP1*

*FOS*

*FOSL2*

*FOXO3*

*GAA*

*GALK1*

*GAPDH*

*GAPDHS*

*GBE1*

*GCK*

*GCNT2*

*GLRX*

*GPC1*

*GPC3*

*GPC4*

*GPI*

*GRHPR*

*GYS1*

*HAS1*

*HDLBP*

*HEXA*

*HK1*

*HK2*

*HMOX1*

*HOXB9*

*HS3ST1*

*HSPA5*

*IDS*

*IER3*

*IGFBP1*

*IGFBP3*

*IL6*

*ILVBL*

*INHA*

*IRS2*

*ISG20*

*JMJD6*

*JUN*

*KDELR3*

*KDM3A*

*KIF5A*

*KLF6*

*KLF7*

*KLHL24*

*LALBA*

*LARGE1*

*LDHA*

*LDHC*

*LOX*

*LXN*

*MAFF*

*MAP3K1*

*MIF*

*MT1E*

*MT2A*

*MXI1*

*MYH9*

*NAGK*

*NCAN*

*NDRG1*

*NDST1*

*NDST2*

*NEDD4L*

*NFIL3*

*NOCT*

*NR3C1*

*P4HA1*

*P4HA2*

*PAM*

*PCK1*

*PDGFB*

*PDK1*

*PDK3*

*PFKFB3*

*PFKL*

*PFKP*

*PGAM2*

*PGF*

*PGK1*

*PGM1*

*PGM2*

*PHKG1*

*PIM1*

*PKLR*

*PKP1*

*PLAC8*

*PLAUR*

*PLIN2*

*PNRC1*

*PPARGC1A*

*PPFIA4*

*PPP1R15A*

*PPP1R3C*

*PRDX5*

*PRKCA*

*PYGM*

*RBPJ*

*RORA*

*RRAGD*

*S100A4*

*SAP30*

*SCARB1*

*SDC2*

*SDC3*

*SDC4*

*SELENBP1*

*SERPINE1*

*SIAH2*

*SLC25A1*

*SLC2A1*

*SLC2A3*

*SLC2A5*

*SLC37A4*

*SLC6A6*

*SRPX*

*STBD1*

*STC1*

*STC2*

*SULT2B1*

*TES*

*TGFB3*

*TGFBI*

*TGM2*

*TIPARP*

*TKTL1*

*TMEM45A*

*TNFAIP3*

*TPBG*

*TPD52*

*TPI1*

*TPST2*

*UGP2*

*VEGFA*

*VHL*

*VLDLR*

*WSB1*

*XPNPEP1*

*ZFP36*

*ZNF292*

**Table S3. Limiting dilution data**

| **A375** | **Cell number of injection** | **Ctrl-sh** | **TIPE-sh** |
| --- | --- | --- | --- |
|  | 10000 | 2/5 | 0/5 |
| **Frequency** | 100000 | 5/5 | 2/5 |
|  | 100000 | 5/5 | 5/5 |

**Table S4. Confidence intervals for 1/ (****stem cell frequency)**

| **A375** | **Lower** | **Estimate** | **Upper** | ***P* value** |
| --- | --- | --- | --- | --- |
| **Ctrl-sh** | 60864 | 18441 | 5588 | *P*<0.001 |
| **TIPE-sh** | 624917 | 202749 | 65780 |  |

**Table S5**	**Primer or siRNA sequences**

| Primer sequences | |
| --- | --- |
| *NANOG* | F: AATACCTCAGCCTCCAGCAGATG  R: TGCGTCACACCATTGCTATTCTTC |
| *NOTCH1* | F: CCTGAGGGCTTCAAAGTGTC  R: CGGAACTTCTTGGTCTCCAG |
| *OCT3/4* | F: CTTGCTGCAGAAGTGGGTGGAGGAA  R: CGGAACTTCTTGGTCTCCAG |
| *SOX2* | F: AAATGGGAGGGGTGCAAAAGAGGAG  R: CAGCTGTCATTTGCTGTGGGTGATG |
| *BMI-1* | F: TGGAGAAGGAATGGTCCACTTC  R: CAGCTGTCATTTGCTGTGGGTGATG |
| *TIPE* | F: TTCAGGCCTCCCTCTTTAACAATC  R: CGTTCGTGGCAGGGGTTATT |
| *HIF-1α* | F: CCAGTTAGGTTCCTTCGATCAGT  R: TTTGAGGACTTGCGCTTTCA |
| *PKM2* | F: TTGCAGCTATTCGAGGAACTCCG  R: CACGATAATGGCCCCACTGC |
| *18S rRNA* | F: CGGCTACCAC ATCCAAGGAA  R: GCTGGAATTACCGCGGCT |
| *NES* | F: AGCTGGCGCACCTCAAGATGT  R: CCTGAAAGCTGAGGGAAGTCT |
| *SOX10* | F: TCATGGTGTGGGCTCAGGCA  R: CGCTTGTCACTTTCGTTCAGC |
| *LDHA(HRE)* | F: TTGGAGGGCAGCACCTTACTTAGA  R: GCCTTAAGTGGAACAGCTATGCTGAC |
| *GLUT1(HRE)* | F: CTGTAATCCCAGCTACTCGG  R: CACGATCTCGGCTCACTGTA |
| *LDHA* | F: ATCTTGACCTACGTGGCTTGGA  R: CCATACAGGCACACTGGAATCTC |
| *GLUT1* | F: CGGGCCAAGAGTGTGCTAAA  R: TGACGATACCGGAGCCAATG |
| siRNA/shRNA sequence | |
| TIPE-sh1 | gtTTCCATCAGGTGGATTATA |
| TIPE-sh2 | ccACCTTAATAGACGACACAA |
| PKM2-sh | Purchased from Santa Cruz Biotechnology, Inc. (#sc-62820-SH) |

**Table S6	Experimental materials**

| Reagent or resource | Source | Identifier |
| --- | --- | --- |
| Antibodies for western blot or immunofluorescence staining | | |
| TIPE | Abcam | ab195810 |
| p-PKM2(Ser37) | Affinity | DF7772 |
| PKM2(Rabbit) | Bioworld | BS6443 |
| PKM2(Mouse) | Invitrogen | MA5-32976 |
| p-PKM2(Tyr105) | Affinity | DF2975 |
| Flag | Bioworld | AP0007 |
| HA | Bioworld | AP0005M |
| His | Zenbio | 350175 |
| CD44, APC | Multisciences | AH04405 |
| LDH | Bioworld | MB63796 |
| HIF-1α(Mouse) | Cloud-Clone | MAA798Hu22 |
| HIF-1α(Rabbit) | Abcam | ab179483 |
| ERK1/2 | CST | #5013 |
| p-ERK1/2 | CST | #4370 |
| Lamin B1 | Proteintech | 12987-1-AP |

**Table S7. The** **clinicopathological characteristics of 48 melanoma specimens**

| **Parameters** | **Number of cases (%)** |
| --- | --- |
| **Gender** |  |
| Female | 25 (52.08%) |
| Male | 23 (47.92%) |
| **Age (years)** |  |
| < 60 | 25 (52.08%) |
| ≥ 60 | 23 (47.92%) |
| **Pathological type** |  |
| Malignant melanoma | 48 (100.00%) |
| **T classification** |  |
| T3 | 4 (8.33%) |
| T4 | 28 (58.33%) |
| Data missing | 16(33.33%) |
| **N classification** |  |
| N0 | 22 (45.83%) |
| N1 | 10 (20.83%) |
| Data missing | 16(33.33%) |
| **Clinical stage** |  |
| II | 27 (56.25%) |
| III | 1 (2.08%) |
| IV | 4 (8.33%) |
| Data missing | 16(33.33%) |
